# Evaluation of TagSeq, a reliable low-cost alternative for RNAseq

**DOI:** 10.1101/036426

**Authors:** Brian K Lohman, Jesse N Weber, Daniel I Bolnick

## Abstract

RNAseq is a relatively new tool for ecological genetics that offers researchers insight into changes in gene expression in response to a myriad of natural or experimental conditions. However, standard RNAseq methods (e.g., Illumina TruSeq^®^ or NEBNext^®^) can be cost prohibitive, especially when study designs require large sample sizes. Consequently, RNAseq is often underused as a method, or is applied to small sample sizes that confer poor statistical power. Low cost RNAseq methods could therefore enable far greater and more powerful applications of transcriptomics in ecological genetics and beyond. Standard mRNAseq is costly partly because one sequences portions of the full length of all transcripts. Such whole-mRNA data is redundant for estimates of relative gene expression. TagSeq is an alternative method that focuses sequencing effort on mRNAs 3-prime end, thereby reducing the necessary sequencing depth per sample, and thus cost. Here we present a revised TagSeq protocol, and compare its performance against NEBNext^®^, the gold-standard whole mRNAseq method. We built both TagSeq and NEBNext^®^ libraries from the same biological samples, each spiked with control RNAs. We found that TagSeq measured the control RNA distribution more accurately than NEBNext^®^, for a fraction of the cost per sample (∽10%). The higher accuracy of TagSeq was particularly apparent for transcripts of moderate to low abundance. Technical replicates of TagSeq libraries are highly correlated, and were correlated with NEBNext^®^ results. Overall, we show that our modified TagSeq protocol is an efficient alternative to traditional whole mRNAseq, offering researchers comparable data at greatly reduced cost.

## Introduction

RNAseq has been widely used to describe differences in gene expression among wild populations, as well as changes in captive or wild individuals' expression following exposure to stimuli (mates, predators, parasites, abiotic stress, toxins). This work has helped uncover the genetic basis of complex traits, implicate genes underlying targets of natural selection, and measure the heritable and environmental components of variation in gene expression [1-6]. However, the most widely used RNAseq protocols are cost-prohibitive for many biologists, including but not limited to researchers in ecological genetics.

Construction of any whole mRNAseq library for the Illumina platform (including Illumina TruSeq^®^ and NEBNext^®^ kits) involves isolating or enriching for mRNA, which is then fragmented and subject to massively parallel sequencing. The resulting data yields sequences for overlapping portions of the entire lengths of the original messenger RNAs (hence 'whole' mRNAseq). This requires high depth of coverage; although sequencing requirements vary depending on sample type, the ENCODE Consortium suggests ∽30 million raw reads per sample as a “best practice” for most RNAseq experiments [7], limiting researchers to a maximum of eight samples per lane of Illumina HiSeq 2500. The high cost of sequencing, combined with high cost of library construction, has forced many studies to use small sample sizes, or pool samples within treatments. This is cause for concern, as meaningful differences in gene expression simply may not be detected with such low-powered sampling designs, and pooled RNAseq may fail to properly account for residual variation in expression.

To resolve problems with whole mRNAseq, several low-cost alternatives have been developed. Most notably, Meyer et al. 2011 [8] presented a 3' Tag-based approach to RNAseq, called TagSeq, that requires little input RNA, involves low library construction costs, and requires many fewer raw reads per sample. By focusing on the 3' end of mRNA fragments, TagSeq reduces the sequencing effort required to characterize a population of mRNAs in a biological sample. This cost-saving does come with some constraints: TagSeq cannot distinguish between alternatively spliced transcripts from a single locus, and will not identify polymorphism or allele-specific expression in much of a genes' coding sequence. However, the benefits of precisely measuring locus-level transcriptional differences with high replication may outweigh the lack of splicing or SNP information for many experiments in ecological systems. However, as presented in Meyer et al. 2011 [8], TagSeq uses a number of outdated methods and enzymes, which may skew the distribution of RNA fragments in the library, with respect to both fragment size and GC content [9]. In addition, the accuracy of TagSeq has not yet been compared to the industry standard TruSeq^®^/NEBNext^®^ which reliably measures moderate and high abundance mRNAs in a sample.

Here, we present a modified protocol intended to increase the accuracy and precision of TagSeq, by incorporating recent findings on polymerase performance, fragmentation methods, and bead-based purification technology into the library construction process. We then tested the accuracy of TagSeq against the industry standard NEBNextM^®^ by sequencing technical replicates of a biological sample, each containing an artificial set of diverse RNAs of known concentration, designed by the External RNA Controls Consortium (hereafter simply “ERCC”).

## Materials and Methods

### Improvements to TagSeq library construction

Briefly, our improved TagSeq library construction method involves 11 steps: 1) isolate total RNA, 2) remove genomic DNA with DNase (if not included in total RNA isolation), 3) fragment total RNA with Mg+ buffer (NEB), 4) synthesize cDNA with a poly-dT oligo, 5) PCR amplify cDNA, 6) purify PCR products with DNA-binding magnetic beads (Agencourt, or made in-house [10]), 7) fluormetrically quantify PCR products (PicoGreen, Life Technologies), 8) normalize among-sample concentrations, 9) add sample-specific barcodes via PCR, 10) pool samples and select a small range of fragment sizes (to maximize output on the Illumina platform) via automated gel extraction (400-500bp, Sage Pippin Prep 2% agarose), 11) quantify concentrations of post-extraction products via Qubit, 12) normalize among pools. This protocol can be completed by a single researcher in three days, and this approach is optimized for 96-well format plates. Improvements over the original protocol are described in Table 1.

**Table 1.** Changes to Meyer et al. 2011. We identified a number of areas where the Meyer protocol could be improved and implemented changes to address these concerns.

| **Problem** | **Solution** |
| --- | --- |
| Quantification of DNA/RNA by spectroscopy is inaccurate. | Fluorescent based quantification with Quant-iT assays. |
| Genomic DNA contamination leads to nonspecific amplification. | Increase DNase treatment to 1.5x concentration at 37°C for 1 hour. |
| Fragmentation of total RNA with Tris buffer produces a wide distribution of fragment sizes. | Precisely fragment total RNA with a specialized Mg<sup>+</sup> buffer. |
| Yield of first strand synthesis is too variable. | Normalize RNA input to 1µg. |
| Variable GC content among fragments can cause dropout of transcripts [9]. | Use AccuPrime Taq polymerase and associated thermal profile for PCR steps [9]. |
| Excessive PCR amplification increases the number of PCR duplicates. | Reduce number of PCR cycles to 12 or less. |
| Purification using solid-phase methods (e.g. spin columns) is not high throughput compatible, inefficient and costly. | Clean with Agencourt AMPure beads, which can be made in-house [10] |
| Post-PCR cDNA amplification yield is highly variable. | Normalize input to 40ng total. |
| Size selection by standard gel extraction is highly variable. | Precise size selection with Pippin Prep automated gel extraction. |
| Mixing individual libraries based on qPCR is slow and expensive. | Normalize lanes of Pippin Prep with Qubit Fluorimeter. |

Total RNA was extracted from six freshly isolated stickleback *(Gasterosteus aculeatus)* head kidneys stored in RNAlater (Ambion). All fish were lab-raised non-gravid females, bred via *in vitro* crosses of wild caught parents. Three fish originated from crosses between parents from Gosling Lake, British Columbia and three fish from crosses between parents from Roberts Lake, British Columbia. Total RNA from all 6 head kidneys was then split, and libraries were constructed with both whole mRNAseq and TagSeq methods. Four whole mRNAseq libraries (NEBNext^®^ directional RNA libraries with poly-A enrichment) were prepared according to the manufacturer' s instruction, by the Genomic Sequencing and Analysis Facility at the University of Texas at Austin, with the addition of ERCC before library construction began, according to the manufacturer's instructions. Whole mRNAsq samples were sequenced on a single lane of Illumina HiSeq 2500 2x100, producing an average of 40.5 million paired-end reads per sample (81 million reads total per sample). Following the addition of ERCC to one technical replicate per biological sample, TagSeq samples were prepared according to Meyer et al 2011 [8], but with changes detailed in Table 1. Four TagSeq samples had two technical replicates (totally independent library builds from total RNA) and a fifth had three technical replicates. TagSeq libraries (29 total, including 17 outside the scope of this work) were sequenced on 3 lanes of Illumina HiSeq 2500 1x100, producing an average of 10.3 million raw reads per sample.

### Bioinformatics

Raw whole mRNAseq reads were trimmed with Cutadapt v 1.3 [11] to remove any adapter contamination. We then mapped the trimmed reads to version 79 of the stickleback genome (with ERCC sequences appended) using BWA-MEM [12], and counted genes using Bedtools [13], producing 20,678 total genes. TagSeq reads were processed according to the iRNAseq pipeline (https://github.com/z0on/tag-based_RNAseq) [14], producing 19,145 total genes.

### Statistical analysis of control transcripts

For each sample, we plotted observed counts of artificial ERCC transcripts against expected values, fitting a simple linear model (observed ∽ expected). We tested for a difference in mean adjusted R^2^ value between library construction methods with a paired t-test (paired by biological sample).

We calculated the Spearman correlation between observed log transformed counts of ERCC transcripts and expected transcript quantity. We tested for a difference in mean Rho values between library construction methods using a paired t-test. We also considered Rho separately for abundance quartiles.

### Statistical analysis of stickleback transcripts

We calculated the Spearman correlation among TagSeq technical replicates. We calculated the Spearman correlation between stickleback head kidney samples which had been prepared using both library construction methods.

### Statistical analysis of inline barcodes

TagSeq, as presented by Meyer et al. and here, uses degenerate inline barcodes on the 5' end of each fragment to identify PCR duplicates. We tested for the random incorporation of these barcodes with a Chi Squared test. We also tested for the effect of increased GC content within each barcode on the number of times that barcode was observed with a Poisson GLM.

## Results

We found that, when fitting a linear model between the expected concentrations of ERCC to observed transcript counts, TagSeq had a significantly higher mean adjusted R^2^ value (R^2^ = 0.89) than NEBNext^®^ (R^2^ = 0.80, Figure 1, observed ∽ expected, paired t-test, t = 18.63, df = 3, p < 0.001). Similarly, the rank correlation between observed and expected ERCC fragments was consistently higher for TagSeq (mean Rho = 0.94) than NEBNext^®^ (mean Rho = 0.87, Figure 2, paired T-test, t = 12.20, df = 3, p = 0.001). TagSeq showed higher mean Rho values for all abundance classes except the third quartile. Most notably, whole mRNAseq performed very poorly in the lowest abundance class (relative concentration of 0.014-0.45 attamols/μl), and TagSeq substantially outperformed whole mRNAseq in the second abundance class (relative concentration of 0.92-7.3 attamols/ul, Figure 3).

**Figure 1.**
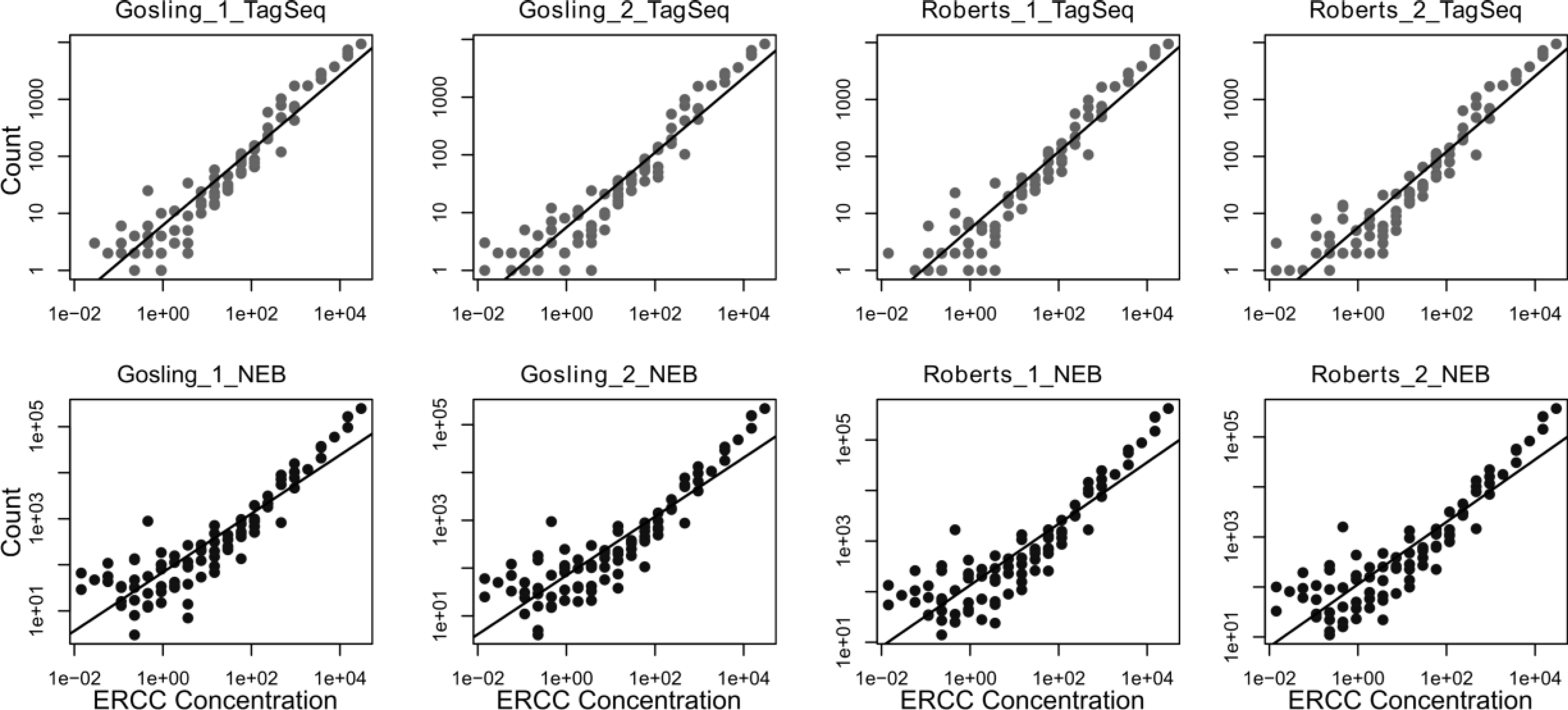
Regression of observed vs. expected ERCC transcripts shows TagSeq has higher adjusted R^2^ values for samples prepared with both methods (paired T-test, t = 18.63, df = 3, p < 0.001).

**Figure 2.**
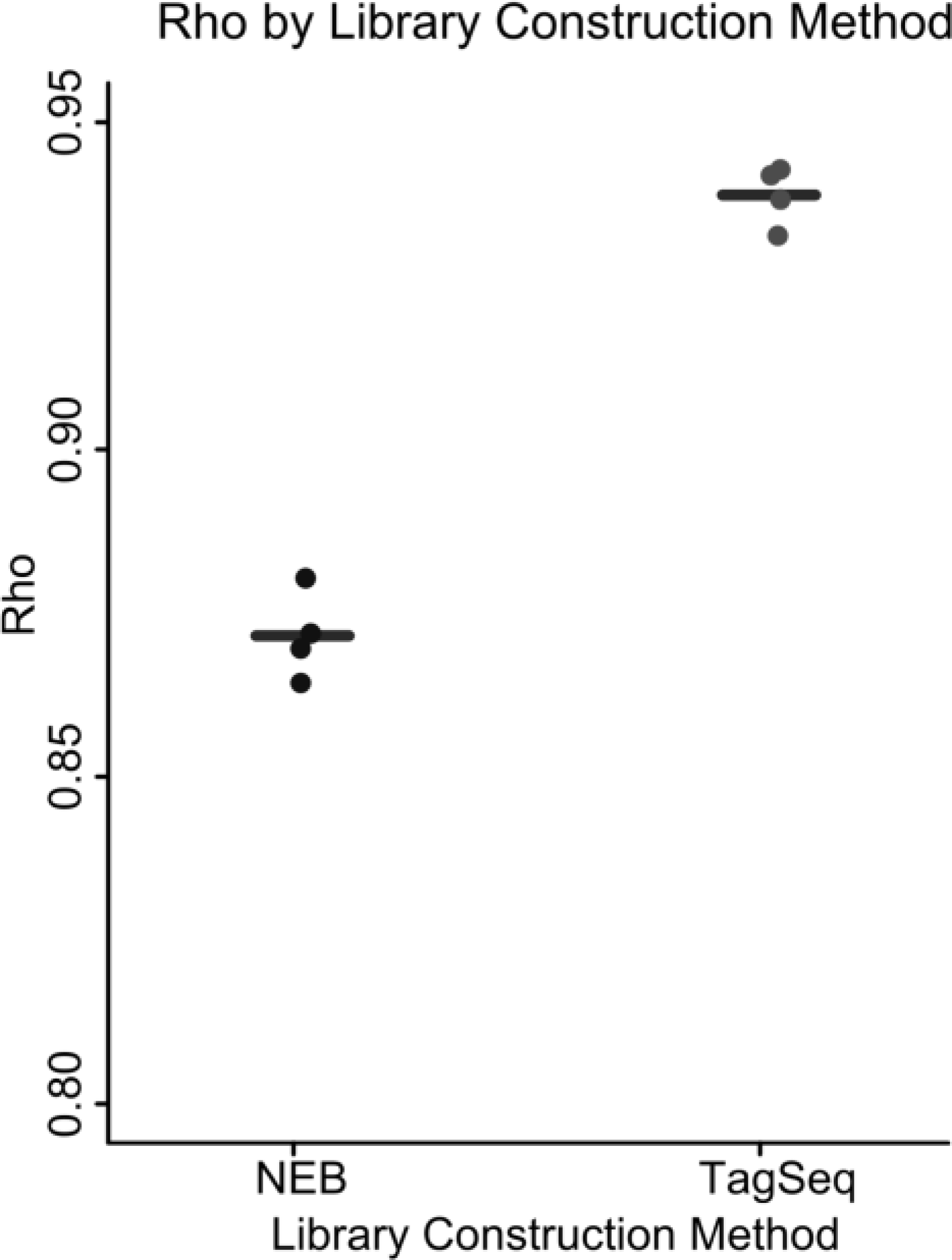
TagSeq more accurately recovers a known distribution of control mRNA fragments (ERCC) than whole mRNAseq (mean Rho for TagSeq is higher than mean Rho for whole mRNAseq, paired T-test, t = 12.20, df = 3, p = 0.001).

**Figure 3.**
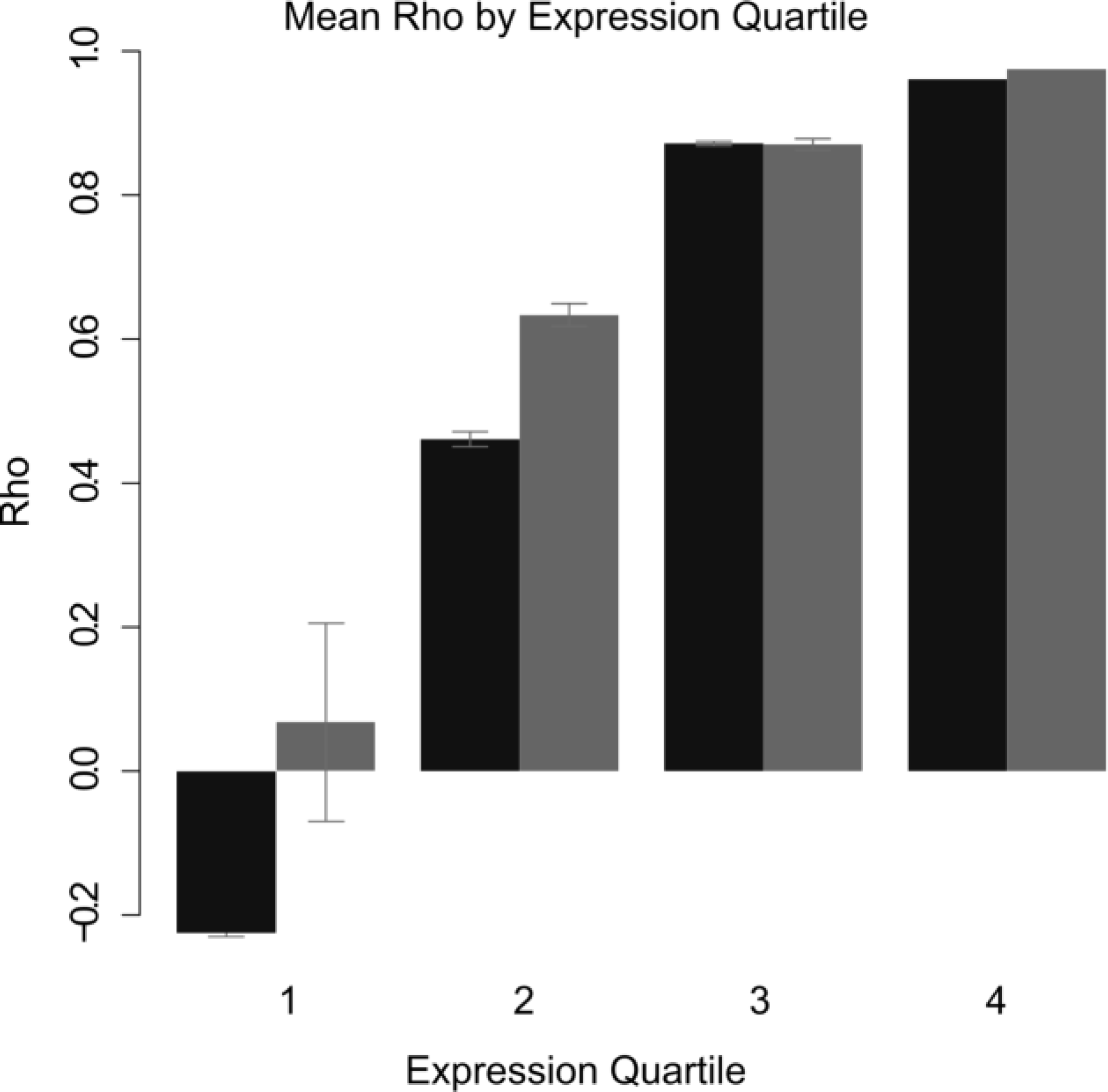
Breakdown of control mRNAs by abundance class shows that TagSeq recovers mRNAs better than TruSeq, especially at lower abundances. Light grey bars are TagSeq, dark grey bars are whole mRNAseq. Fences indicate standard error.

With respect to stickleback (non-control) sequences, the mean Rho among technical replicates of TagSeq samples was 0.96 (n=5, calculate Rho for each biological sample and average). Due to the high cost of NEBNext^®^ library generation and sequencing (∽$340 per sample), we did not perform technical replicates using this method. We found a strong significant positive correlation between stickleback gene counts generated with TagSeq and whole mRNAseq (Rho = 0.74, p < 0.001). This is likely and underestimate of the actual correlation between the two library construction methods because whole mRNAseq performs very poorly when RNAs are in moderate to low abundance (first and second abundance classes, Figure 3). Given that 9,572 loci are in the bottom half of gene counts, even small differences in absolute counts between the methods will strongly influence the rank-based statistic.

We also wished to compare our new method with that of the original TagSeq protocol, but cannot make a direct comparison with the available samples. Meyer et al. (2011) evaluated their accuracy by comparing fold-differences in differentially expressed genes (between experimental treatments), whereas we measured accuracy using the estimates of relative abundance of ERCC. Keeping in mind these different benchmarking methods, we can draw a rough comparison. The original TagSeq method yielded a correlation of r = 0.86 between TagSeq estimates of fold-change expression, and qPCR measures of the same fold change (a “known” benchmark). In contrast, our protocol yields a correlation of Rho = 0.94 between our relative abundance estimates, and the known ERCC relative abundances. We infer that the new protocol performs at least as well, and probably better, than the previous protocol, at generating expression level estimates that resemble known values.

The iRNAseq pipeline includes the removal of PCR duplicates, which are a common problem in many library construction methods [9]. Any reads which meet two criteria are called PCR duplicates and removed: 1) identical in-line barcodes; four degenerate bases at the start of each read, and 2) the first 30 bases of sequence after the in-line barcode are identical. The removal of PCR duplicates substantially reduces the number of TagSeq reads in each library (mean reduction of 70.3%, n = 12). However, this avoids potential bias introduced by PCR, namely over representation of smaller fragments. We found that inline barcodes were incorporated non-randomly (Chi Square = 10,500,000, df = 63, p ≪ 0.001). We found that increased GC content in the inline barcode significantly reduced the number of times a barcode was observed. For every G or C added to the inline barcode, the expected value of the number of observed barcodes is reduced by ∽2.9% (count ∽ gcContent, family = poisson, Pg cContent -0.133, p < 0.001).

## Discussion

We present a number of methodological improvements to the TagSeq method of Meyer et al. 2011, and taken the important next step of comparing the new protocol to the NEBNext^®^ kit, the industry-standard for whole mRNAseq. Overall, our results illustrate that the updated TagSeq method offers researchers the ability to dramatically increase sample sizes in gene expression analyses, which will facilitate testing for more subtle transcriptional differences than traditional whole mRNAseq methods.

While TagSeq has been used predominantly in corals [14, 15], it should be applicable to nearly all metazoans. However, we caution researchers to perform several basic checks during TagSeq library construction, most especially ensuring the size distribution of RNA fragments is as narrow as possible during total RNA fragmentation. We recommend evaluating the results of various total RNA fragmentation times via BioAnalyzer. Fragments should be larger than 100 bp and smaller than 500 bp (see supplementary materials). Here we were interested in evaluating the robustness of the TagSeq method for threespine stickleback, and therefore sequenced stickleback transcripts more deeply than required for an accurate estimate of gene expression across the majority of expressed loci (we generated an average of 10.3 million raw reads per sample). We recommend that researchers aim for ∽5-6 M raw reads per sample if the goal is to measure the top 75% of all expressed mRNAs in a sample, as this has produce sufficient gene counts for robust statistical power in an invertebrate, a plant, and stickleback (M. Matz and T. Jeunger, personal communications).

In this project, we intentionally under-loaded our TagSeq libraries on the HiSeq lane by 15% (0.0017 pmols loaded), anticipating that low base diversity on the 5' end of the fragments (the inline barcode used to detect and remove PCR duplicates) would lead to poor clustering. However, quality metrics from the HiSeq run indicate that this is not a problem. We observed ∽500-600 clusters per mm^2^ on each tile, and the majority of these clusters passed filtering (low base diversity or overclustering would generate large numbers of clusters with few passing filtering). We therefore recommend that users load the expected quantity or even 10-20% extra material on each lane of HiSeq (see supplementary material). Overloading TagSeq libraries may help to increase raw read yield, relative to NEBNext^®^ (optimally clustered at ∽1000 clusters per mm^2^ when 0.002 pmols loaded). We also emphasize that small fragments need to be removed from TagSeq libraries, as they will more easily cluster on the HiSeq, reducing read output. These may be identified by BioAnalyzer and removed with additional bead clean-ups.

Several of our methodological changes aimed to mitigate the number of PCR duplicates, which are artifacts of all PCR-related methods. First, we predicted that adding two additional degenerate bases to the inline barcodes (the first four sequenced bases of every adapter, which were coded as degenerate bases in the old TagSeq method) would not only increase our ability to detect independent transcripts from PCR duplicates, and also increase base diversity on the 5' end of fragments, thereby increasing the number of clusters passing Illumina's quality filters. However, this alteration did not ameliorate the problem of PCR duplicates or increase the number of raw reads generated in each lane (data not shown). In the future we recommend that protocol users consider replacing the degenerate bases in inline barcodes with 3-nitropyrrole, as this should better randomize which bases are incorporated during initial round of PCR [16]. Second, we limited our number of PCR cycles to 12. Empirically testing the effects of PCR cycle number on TagSeq accuracy was outside the scope of the present study. However, it is widely accepted that the best way to limit bias is to reduce the number of PCR cycles during cDNA amplification as much as possible [9].

In summary, we show that the improved TagSeq method has both benefits and drawbacks compared to traditional whole mRNA sequencing. While our TagSeq libraries did not generate optimal numbers of clusters on the HiSeq platforms, but we identify several potential solutions to the problem. Regardless of the slightly lower number of raw reads, our improved TagSeq method remains far and away much more cost effective than whole mRNAseq. At maximal efficiency (32 individuals per sequencing lane), our method was able to produce highly accurate, transcriptome-wide gene counts for only ∽$33 per sample, including sequencing costs (one lane of HiSeq 2500 V3 chemistry with ∽5.6 M raw reads per sample). This low cost and high reliability offer molecular ecologists the opportunity to vastly increase sample sizes and increase replication to uncover new patterns in gene expression.

## Acknowledgements

All live animal research was approved by the UT Austin IACUC [protocol AUP-2013-00012]; and collections were approved by the British Columbia Ministry of Environment [Scientific Fish Collection permit NA09-52421]. We wish to thank Mikhail Matz, Marie Strader, and the Juenger lab for fruitful discussion on improvements to the TagSeq method both during library construction and analysis. Figures 2 and 3 were generated using plotting functions written by Luke Reding (https://github.com/lukereding/redingPlot). This work was supported by the Howard Hughes Medical Institute (DIB).

## Data Accessibility

Meta data, code for raw read processing, gene counts, code for statistical analysis, and plotting of data, BioAnalyzer .XAD files, and HiSeq quality metrics, and detailed protocol are located in DRYAD entry: http://dx.doi.org/10.5061/dryad.vq275

Raw sequence reads will be stored on Corral, a permanent data repository with multiple, independent backups, located and owned by the University of Texas at Austin Texas Advanced Computing Center. Users can download data at any time via secure copy.

## Author Contributions

BKL, JNW, and DIB jointly designed the research. BKL carried out all library construction improvements. BKL, and JNW analyzed data. BKL wrote the manuscript, with comments from JNW and DIB. All authors approved the final version.

